# A Reproducible and Extensible Benchmark of Supervised Cell Type Annotation Tools for Cytometry Data

**DOI:** 10.64898/2026.06.02.729500

**Authors:** Frederik Kirk, Andrea Sonnenholzner, Helene Scheel Wegener, Javier Herranz del Cerro, Signe Modvig, Lars Rønn Olsen

## Abstract

High-dimensional cytometry technologies such as flow cytometry (FCM) and mass cytometry (CyTOF) are central to immunophenotyping in research and clinical practice. While manual gating remains the standard for cell population annotation, it is time-consuming, difficult to scale, and subject to inter-operator variability. Supervised annotation methods have emerged as a way of scaling manual annotation work, yet independent benchmarks for comparing these tools remain limited and quickly become outdated.

This study presents a reproducible and extensible benchmark of supervised cytometry annotation tools implemented within the OmniBenchmark framework. Five supervised annotation methods were evaluated, spanning linear models, nearest-neighbor approaches, tree-based classifiers, mixture-rule systems, and deep learning, across eight publicly available datasets carefully selected to cover technologies, tissues, panel designs, and healthy and disease contexts. Using a sample-centric cross-validation design that reflects common reference-mapping scenarios, overall and per-population F1 scores, performance on rare populations, runtime, and robustness to reduced training set sizes was tested.

Performance varied substantially across datasets and was not fully explained by dataset size or dimensionality, highlighting both operator dependence in annotation and the importance of biological context, cohort heterogeneity, and population imbalance. Less prevalent populations (<1%) remained a key challenge for most methods. Downsampling analyses showed that moderate reference sizes were often sufficient to achieve near-maximum performance.

Rather than ranking methods, this benchmark provides a standardized and transparent framework for evaluating annotation tools under realistic deployment conditions. As a living resource, the OmniBenchmark implementation supports continuous integration of new datasets, tools, and metrics for both tool developers and end users annotating datasets. This enables ongoing, reproducible method comparison and informed tool selection for diverse cytometry applications.

## Introduction

Cytometry comprises a family of high-throughput, single-cell measurement technologies that quantify cellular properties and marker expression to identify biologically meaningful cell populations [1,2]. These assays are central to immunophenotyping in basic research and clinical practice, including disease diagnosis and monitoring, as well as assessment of responses to therapy [1] or vaccination [3].

Commonly used platforms for immunophenotyping are fluorescence-based flow cytometry (FCM) and mass cytometry (CyTOF)[4]. FCM relies on panels of fluorophore-conjugated antibodies and optical detection to quantify marker expression [1,4]. CyTOF combines antibody-based labeling with elemental mass spectrometry, where antibodies are tagged with heavy metal isotopes and quantified by time-of-flight detection, enabling higher-parameter panels with reduced signal overlap between channels [2,4,5], and reduced need for spillover compensation controls [2,4]. These advantages come with trade-offs: compared with FCM, CyTOF typically acquires fewer cells per unit time [4,5] and requires dedicated instrumentation and technical expertise, which can limit accessibility outside specialized core facilities [5].

Regardless of platform, translating high dimensional marker measurements into discrete, biologically meaningful cell populations remains challenging. Conventionally, annotation is done via manual gating, in which analysts iteratively draw boundaries on one or two dimensional marker projections to define populations using hierarchical strategies [1,4,6]. While manual gating remains the common practice, it is recognized as a bottleneck in large studies due to limited reproducibility and scalability [7]. Reported gates can vary between expert analysts and across centers, contributing to inter-operator variability [7,8]. Moreover, manual gating is time consuming, and the burden increases with higher-dimensional panels [8,9].

These limitations have motivated the development of computational approaches for unsupervised clustering and, more recently, automatic annotation methods that assign identity and state labels to cells with minimal expert intervention [7]. The tools evaluated in this study reflect the diversity of current methods spanning from linear classifiers (CyTOF Linear Classifier [10]) and nearest-neighbor based classification (CyAnno [11]), to tree-based approaches (CyGATE [12]), mixture-model and rule-based methods (GateMeClass [13]), and deep learning-based models (DGCyTOF [14]). These tools differ in their handling of unlabeled cells, scalability to large datasets, and requirements for training data.

Despite this progress, selecting an appropriate annotation method for a given study remains challenging. Most comparative evaluations of supervised auto-gating methods are embedded in papers that introduce a new tool, hence, evaluation choices are inevitably entangled with development. Truly independent, third-party benchmarks designed primarily to compare existing annotation methods rather than to present a new one remain scarce [15], and existing ones focus on unsupervised population discovery (clustering) rather than supervised label assignment [7]. A recent study partially closes this gap by benchmarking manual gating, 23 clustering tools, and four supervised auto-gating methods under a unified pipeline [16]. However, with only one of the tools evaluated in this work being included, this comparatively comprehensive effort highlights how quickly evaluations can become outdated. Further, because only a minority of mass cytometry datasets are publicly accessible, dataset reuse is concentrated in a small number of widely used studies. In some instances, these datasets have become de facto standards for both development and evaluation, raising the risk of data leakage and over-optimistic performance estimates that do not generalize beyond a given benchmark [15].

In this work, we present a reproducible and extensible benchmark of supervised cytometry annotation tools, designed to support both method developers and end users. We evaluate five currently implementable tools across eight diverse datasets spanning both flow and mass cytometry platforms, reporting accuracy, sensitivity to rare populations, and computational scalability within a standardized pipeline. A gold standard benchmark requires consistent preprocessing, explicit evaluation criteria, and transparency in method implementation principles, which we formalize using the omnibenchmark workflow system [17]. Omnibenchmark formalizes benchmarks as version controlled, modular pipelines, enabling continuous integration of new datasets, tools, and metrics while preserving reproducibility. Unlike traditional static benchmarks, this framework functions as a living resource where users can:

1. execute locally to evaluate how methods perform on their own datasets prior to committing to a particular approach, and,
2. contribute new data, methods, and metrics, ensuring fair and comprehensive comparisons dynamically beyond this work.

This design ensures the benchmark remains up-to-date and scientifically relevant as the field evolves, addressing the problem of benchmark staleness while maintaining the rigor needed for method comparison. The goal of this benchmark is not to rank methods from best to worst performing, but rather to objectively report their performance across diverse experimental contexts and practical constraints. Accordingly, we quantify how performance varies across platforms, tissues, and panel designs, with attention to identification of rare populations.

## Methods

### Data

To benchmark cell type annotation tools across FCM and CyTOF data, we assembled eight publicly available datasets spanning different sample sources and biological contexts. We include two datasets widely used in comparison analyses: the ‘Levine’ (HumanBoneMarrow_Cyt) dataset [15], representing human bone marrow, and the ‘Samusik’ (MouseBoneMarrow_Cyt) dataset [18], derived from mouse bone marrow. These were complemented by the ‘Bodenmiller’ (PBMC_Cyt) dataset of healthy human PBMCs [19]. This inclusion allows for the assessment of algorithmic robustness against variable pre-processing and data distribution types. To evaluate the resolution of dynamic immune states, we included the ‘Rybakowska’ (StimBlood_Cyt) dataset [20], which challenges tools to distinguish between resting and stimulated immune populations in whole blood.

To serve as the standard reference for conventional fluorescence-based FCM in healthy tissue, we selected the ‘FlowCyt’ (HumanBoneMarrow_flow) dataset [21]. Furthermore, to test annotation accuracy under conditions of severe immune dysregulation, we incorporated two disease-specific studies. We utilized the COVID-19 Immune Phenotyping study [22], stratified into Healthy (PBMC_flow) and covidPBMC_flow cohorts, to measure performance shifts between baseline homeostasis and severe infection. Similarly, to extend this pathological benchmarking to mass cytometry, we utilized the Chikungunya virus infection dataset (ChikVirusPBMC_Cyt) [23], which profiles complex myeloid and lymphoid reorganizations during acute viral response. All datasets were standardized and processed within the omnibenchmark framework [17], with expert manual gating provided in the original studies serving as the ground truth for performance evaluation. Dataset characteristics, including platform, sample counts, marker panel size, number of labeled populations, and transformation status, are summarized in **Table 1**.

**Table 1:** Overview of the datasets used during this study. **Characteristics of the mass and flow cytometry benchmarking datasets.** Summary of the eight datasets utilized to evaluate the automated cell type annotation tools. The table details the biological context (tissue source, species, and healthy/disease state), experimental platform (CyTOF or FCM), and overall dataset dimensionality. Dimensionality metrics include the number of biological samples, the number of quantified markers, and the total number of expertly annotated cell populations (ground truth). The dataset abbreviations provided are used consistently throughout the manuscript and figures.

| Datas<br>et | Descripti<br>on | Platfo<br>rm | N<br>sampl<br>es | N<br>marke<br>rs | N<br>populati<br>ons | N<br>batches | N cells<br>total | Abbreviation<br>s | Referen<br>ce |
| --- | --- | --- | --- | --- | --- | --- | --- | --- | --- |
| FR-FC<br>M-Z3Y<br>R | Healthy +<br>Stim<br>Blood | CyTO<br>F | 32 | 38 | 28 |  | 15,631,02<br>2 | StimBlood_Cyt | [20] |
| Boden<br>Miller<br>XL | Healthy<br>PBMC | CyTO<br>F | 16 | 33 | 8 |  | 163,036 | PBMC_Cyt | [19] |
| FlowCyt | Healthy<br>Bone<br>Marrow | FCM | 30 | 12 | 5 |  | 2,059,816 | HumanBoneMar<br>row_flow | [21] |
| Samusi<br>k | Bone<br>Marrow<br>Mouse | CyTO<br>F | 10 | 39 | 15 |  | 472,988 | MouseBoneMarr<br>ow_Cyt | [18] |
| Levine | Bone<br>Marrow<br>Human | CyTO<br>F | 2 | 32 | 15 |  | 72,570 | HumanBoneMar<br>row_Cyt | [24] |
| FR-FC<br>M-Z2K<br>P | Human<br>COVID-1<br>9 Immune | FCM | 6 | 24 | 6 |  | 3,564,728 | covidPBMC_flo<br>w | [22] |
|  | Phenotyping<br>(PBMCs) |  |  |  |  |  |  |  |  |
| FR-FC<br>M-Z2K<br>P | Human<br>Healthy<br>Immune<br>Phenotyping<br>(PBMCs) | FCM | 43 | 24 | 6 |  | 1,352,280 | PBMC_flow | [22] |
| FR-FC<br>M-Z238 | Immune<br>profiling<br>of<br>chikungunya virus<br>infection | CyTOF | 43 | 37[23] | 16 |  | 2,410,903 | ChikVirusPBMC<br>_Cyt | [23] |

### Data preprocessing

In order to benchmark models under a common data representation, we standardized all datasets to one CSV per biological sample, where each row corresponds to a cell and each column corresponds to a retained marker/channel; the last column, *label*, stores the population annotation. We performed dataset-agnostic formatting steps only (no additional gating beyond what was provided by the dataset authors): channel names were normalized to resolve naming inconsistencies across files, and non-informative acquisition channels were removed. Specifically, we excluded instrument-only/time-of-flight–related channels (e.g., *TIME, SS PEAK, SS TOF*) and excluded cell-viability channels when present. The remaining columns correspond to the marker set used for cell-type discrimination in each dataset.

For FlowRepository datasets, annotations were obtained by applying the published gating strategy stored in the accompanying FlowJo workspace files (.wsp; e.g., FR-FCM-Z3YR). For FCM datasets, we generated per-case CSVs from the raw FCS population files and produced (i) a 6-class full dataset including an ungated class while retaining the 12 channels used in the original study. The StimBlood_Cyt, HumanBoneMarrow_Cyt, and MouseBoneMarrow_Cyt reference datasets were obtained from HDCytoData and used with the accompanying labels.

Due to performance load on the original host of the datasets (flow-repository [25]), the selected datasets were extracted and mirrored on a separate repository [26], since the omnibenchmark requires redownloading datasets at every run to ensure no leakage between runs. Each dataset was compressed due to file-size limitations and easier handling using zstandard [27].

See this github repository for the complete list of files [26].

### Tools

For this benchmark we included tools developed for automated cell annotation in cytometry data that can be executed end-to-end with no user intervention or intermediate decisions during a run, enabling fair comparisons under a standardized benchmarking pipeline. Tools that required intermediate user decisions or were incompatible with the Omnibenchmark framework were excluded.. A custom wrapper function was used for parsing inputs and reporting results. The included tools, along with their implementation details, are summarized in Table 2. The tools span a range of algorithmic approaches and were evaluated at the most recent commits to ensure reproducibility.

**Table 2:**
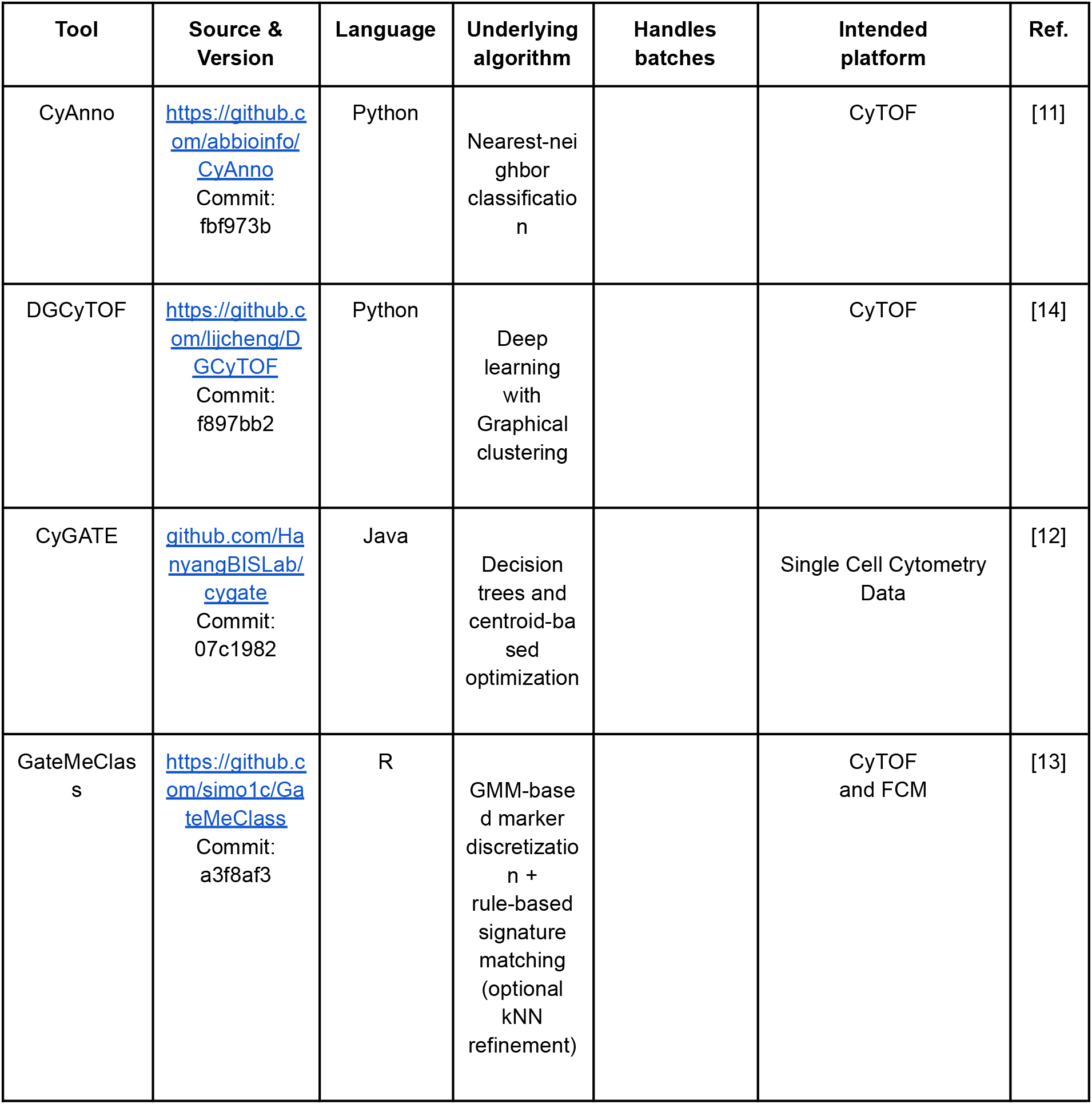
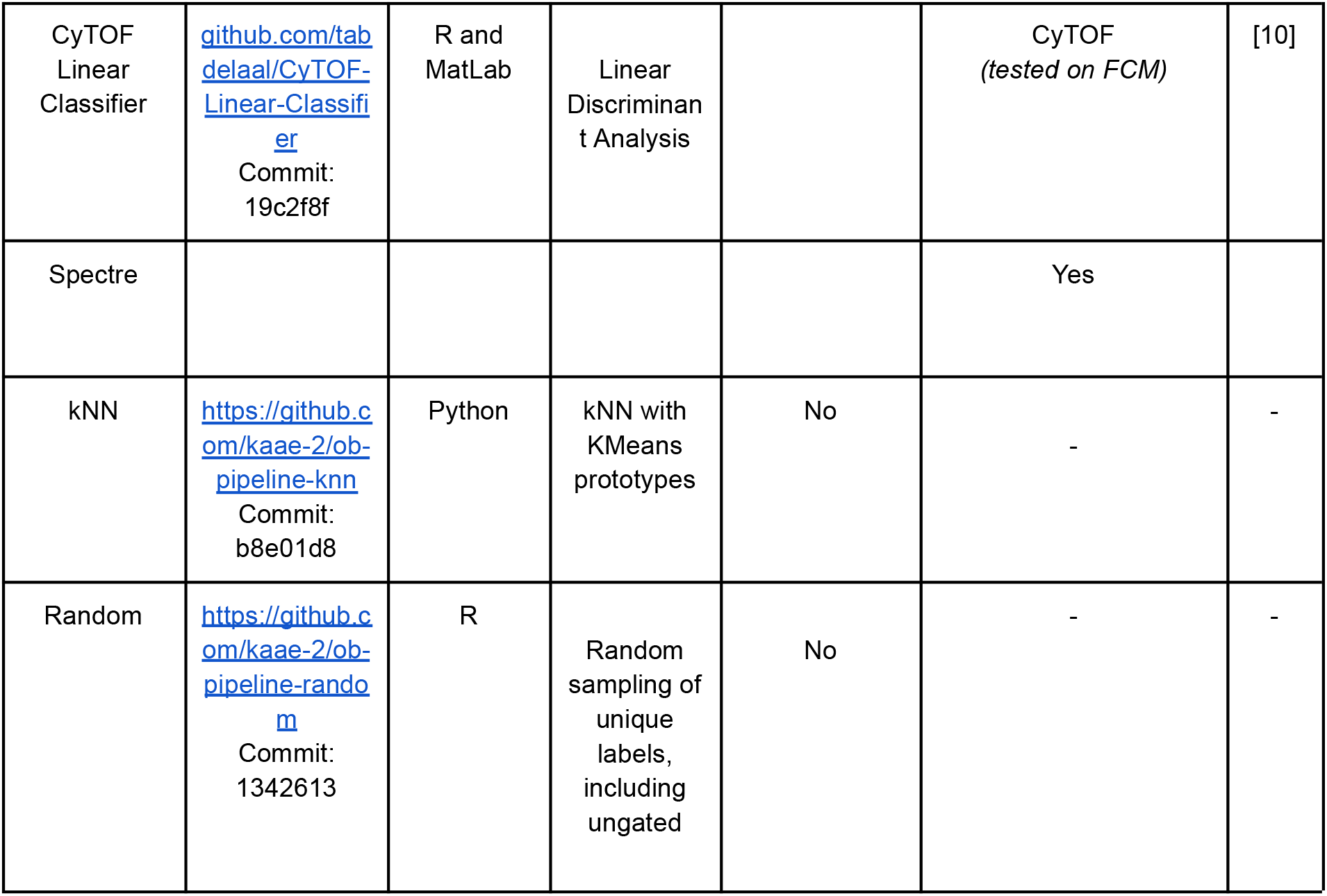
Supervised cell type annotation tools included in this study. Overview of the computational methods evaluated in this benchmark. For each tool, the specific source repository and commit hash are provided to ensure exact reproducibility. The table details the primary programming language(s) utilized, a brief description of the core underlying algorithm, the original cytometry platform the tool was intended for or validated on, and the corresponding literature reference. The kNN and Random methods serve as standard performance baselines for the evaluation.

### CyAnno

CyAnno utilizes kernel density estimation (KDE) on principal components to isolate “landmark” cells that represent the core of a population. These landmarks form a cell-type-specific training set that includes both target events and “ungated” background noise. This benchmark specifically employed the XGBoost implementation to train binary classifiers. Final assignments are based on posterior probabilities; any cell failing to reach a 0.5 threshold is labeled “Unknown,” allowing the model to reject cells that do not fit the reference atlas. [11].

### DGCyTOF

DGCyTOF uses a deep feedforward neural network (three hidden layers with 128, 64, and 32 nodes respectively) with ReLU activation and a Softmax output. Cells with low prediction probabilities are initially labeled “unknown” and in the full workflow, these cells are then clustered using UMAP and HDBSCAN to determine if they represent novel biological populations or simply technical artifacts [14]. For this benchmark, we focused on the classification component, utilizing the deep learning module to transfer labels from the reference to the query datasets while assessing its ability to flag unassigned cells.

### CyGATE

CyGATE is a semi-automated classification tool that combines decision trees with centroid-based optimization. It first identifies marker-specific boundaries to define cell-type centroids. During classification, test cells are tentatively labeled via the decision trees and then refined through centroid optimization. This iterative process ensures the model adapts to the specific distribution of the test sample. Cells that fail to meet the optimized confidence criteria are tagged as “ungated.”[12]

### GateMeClass

GateMeClass allows supervised and semi-supervised annotation of flow-cytometry data [13]. For this benchmark we evaluate the semi-supervised method which consists of classifying marker intensities in “low”, “medium” and “high” states using per-marker Gaussian mixture models. For this benchmark, we evaluated the two available configurations: Equal (E), which assumes identical Gaussian distributions across markers, and Variable (V), which optimizes distributions individually. This allows for a rule-based matching of cell signatures against the GMM-defined states. Notably, the GateMeClass scales linearly with both training set size and testing set size due to need for pairwise comparisons. For this reason, GateMeClass was excluded for larger datasets.

### CyTOF Linear Classifier (LC)

CyTOF Linear Classifier is a supervised learning tool based on Linear Discriminant Analysis (LDA). It calculates a linear combination of markers to maximize the distance between populations in a multi-dimensional space. Following projection into the LDA space, a confidence score is generated for each cell. We applied the default 0.7 (70%) threshold, classifying any cell falling below this mark as “Unknown.”[10]

### k-Nearest Neighbor (kNN)

To establish a performance baseline, we included a simple kNN classifier. In this implementation, K is dynamically selected based on the number of populations, *L*_*pop*_ and the size of the smallest gated cluster: *K* = *min* (2 · *L*_*pop*_, *N*_*min*_*pop*_). To improve scalability and prevent 1-to-1 comparison overhead, we used MiniBatchKMeans to generate eight prototypes per class, effectively summarizing the training data without losing population variance.

### Random Classifier

As a negative control, this classifier assigns labels by randomly sampling from the distribution of the training set. By ignoring marker expression profiles entirely, it provides a lower bound of performance, helping to quantify how much of a tool’s accuracy is due to the algorithm versus the underlying frequency of the populations.

### Metrics

For each dataset, tool, and cross-validation fold, we compared predicted labels to the published manual gating labels and computed F1 scores at two levels: an overall F1 summary per dataset–fold and per-population F1 scores across test samples. For per-population analyses, we also report a support-weighted F1 score, computed as the mean of per-population F1 values weighted by the number of true cells in each population,so that more abundant populations contribute proportionally to the summary. F1 is defined as 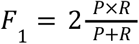, where 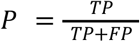, and 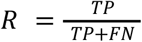. These measures provide a balanced view of both general and population specific performance. Furthermore, the runtime was recorded for each cross-validation to compare efficiency of analysis.

### Implementation

We implemented the benchmark using the OmniBenchmark framework [17], which structures evaluations as a modular pipeline separating data import, method execution, and metric calculation. Each stage is containerized and version controlled, and the full workflow is pinned to specific git commit hashes [28] to ensure results are reproducible from an exact pipeline state. Continuous integration is used to rerun the pipeline under the same software environments when components change. This design also allows new datasets or methods to be added without altering the overall workflow structure.

Since omnibenchmark only specifies that stages are implemented as command-line interface tools, each step in the benchmark was configured as follows:.

- **Environment set-up**. For each step in the pipeline, an environment (containing the packages and runners needed for that step), was created.
- **Data import**. Datasets were downloaded from the repository [26], extracted and loaded into csv files into datasets, extracting meta-data from the datasets. Metadata includes file size, number of samples, number of cells, dataset name for post processing at future stages. Furthermore, a random order of the samples was created (using a fixed seed for reproducibility), to be used for preprocessing into training-test sets.
- **Preprocessing**. The data was split into 5 separate cross-validation sets. For the training-set split, 1 training sample was fully labelled, all other samples were test samples to simulate a realistic “reference mapping” scenario where annotated data is limited to a single biological source. Labels were cast as integers for later processing, and all labels indicating ungated cells, such as debris or N/A values, were cast as the same label (0) for later handling. For datasets with fewer than 5 samples (Levine), a number of cross-validations equal to the sample number was performed. Headers were stripped from the datasets and labels in order to simplify model processing. To simulate lower data-resolution and test performance of models’ boundary abilities, a dataset was subsampled. Training sets and labels were proportionally reduced to 1k,5k,10k,50k, and 100k cells. The testing sets remained full-size. All output data were then compressed to a tar.gz archive into single files for easier handling at the following step.
- **Analysis**. Each model was run using a wrapper function that extracted relevant input parameters, such as path to data, model name and expected output folder, Afterwards, data was extracted from tar archives, before any model specific preprocessing such as filtering ungated values, transforming the data to different scale, and/or transposing data. Then, each model performed training using training data and labels, and all test data was annotated for each test sample into a prediction label vector. Finally, all the prediction labels were compressed to a tar.gz archive for downstream processing. Each model ran independently on each dataset-crossvalidation and produced a set of predictions for each test sample. During testing, GateMeClass performance was found to be slow, (>4 hours per cross-validation run), on the larger datasets and was therefore manually excluded from running on those.
- **Metrics**. F1 scores were calculated for each cross-validation fold as described in *metrics*.
- **Collector**. The computed metrics were collected to extract for plotting.

All runs were performed on a workstation with an AMD Ryzen 9 6900HX CPU and 64 GB RAM running Arch Linux 2025.11.01, using up to 12 cores for parallel execution. The complete configuration (including pipeline commit hashes and container definitions) is recorded in the benchmark repository [29].

## Results

We implemented the tool evaluation as a fully runnable benchmark within the OmniBenchmark framework. The pipeline standardized inputs and outputs across tools, applied a sample-centric reference-mapping evaluation design, and aggregated performance and runtime metrics into analysis-ready tables. Because OmniBenchmark is designed as a living benchmark, the workflow can be rerun as new tools or datasets are added, supporting continuous, standardized re-evaluation. Figure 1 shows a visual representation of the benchmark topology.

**Fig. 1.**
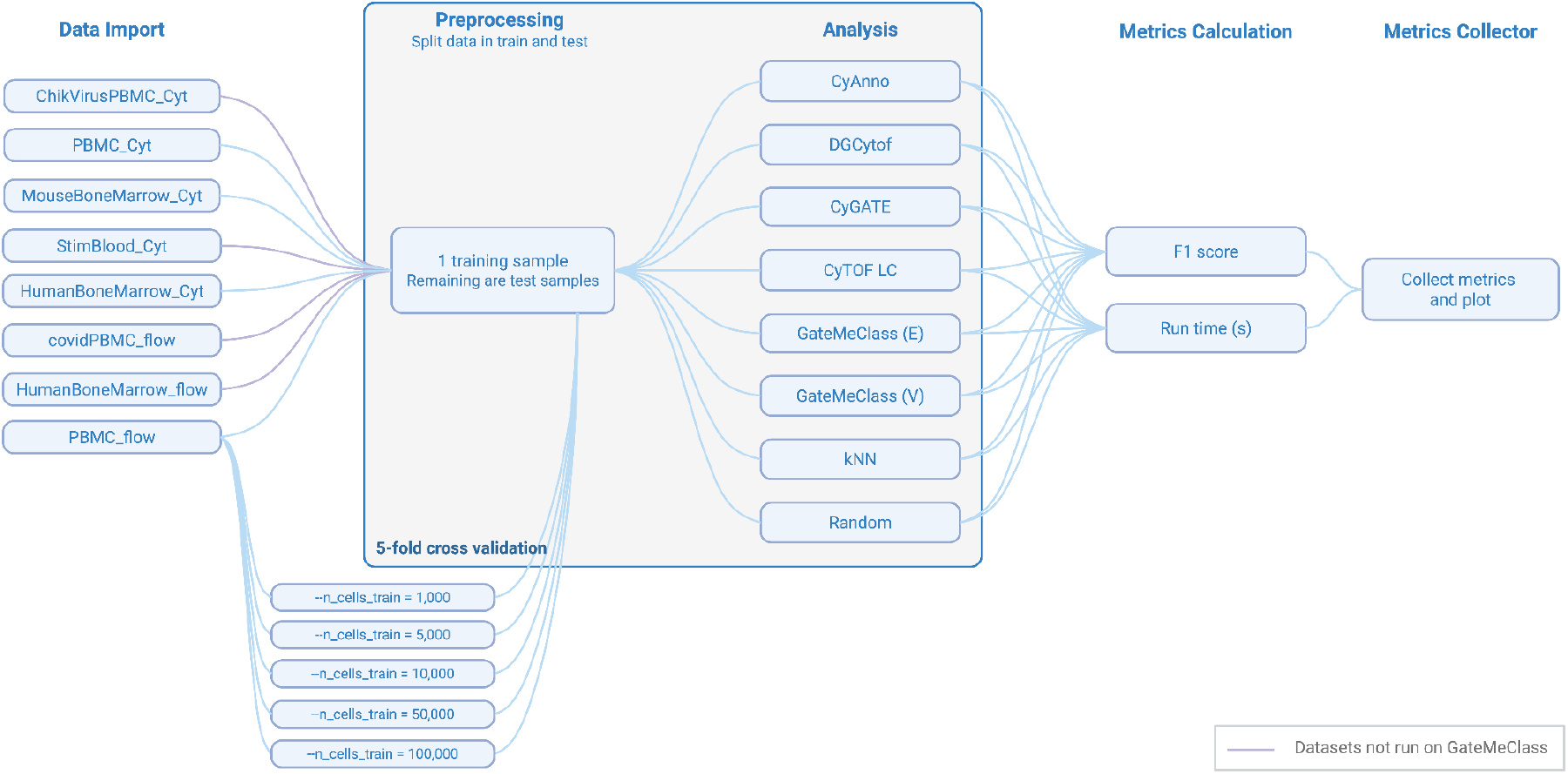
OmniBenchmark topology for the cytometry annotation benchmark. The benchmark is implemented as a modular workflow with five stages: Data Import, Preprocessing, Analysis, Metrics Calculation, and Collection. Datasets are imported and standardized, then transformed into reference/query splits using a sample-centric reference-mapping design with 5-fold cross-validation; a parallel branch performs reference-size downsampling (1k–100k training cells). Tool-specific wrappers execute each method and write predictions in a shared output format, enabling tool-agnostic computation of F1 and runtime and aggregation for visualization. Purple connections denote dataset-tool combinations not evaluated due to runtime constraints.

### Performance evaluation

For a comparison overview of the evaluated tools, Figure 2 summarizes overall annotation accuracy. We report mean macro-F1 for each tool–dataset pair under the sample-centric cross-validation design, alongside distributions across folds and datasets to highlight both average performance and variability. This provides a compact view of robustness across platforms and biological contexts.

**Fig. 2.**
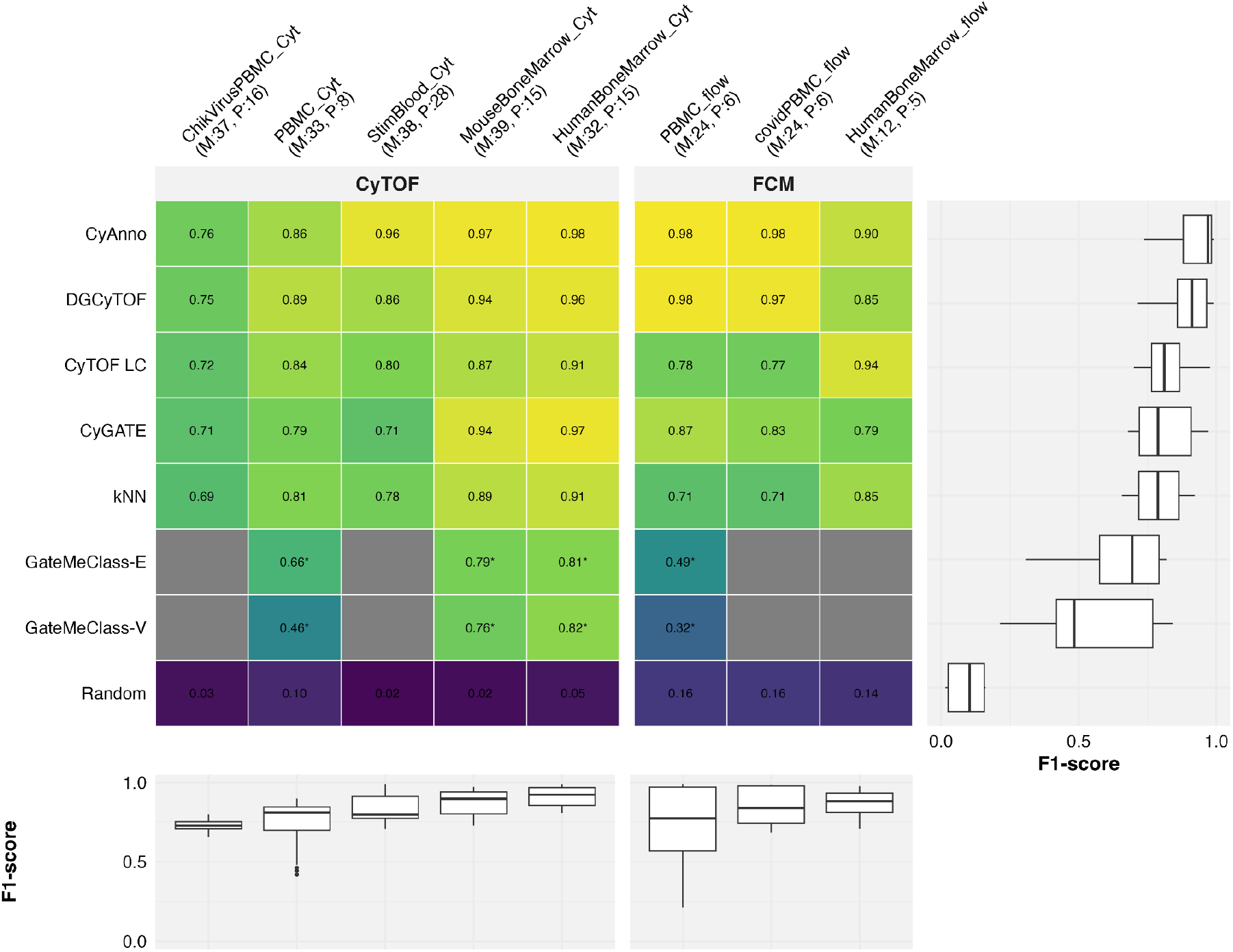
Classification performance across CyTOF and FCM benchmarking datasets. Heatmap showing the mean macro F1 scores of the automated cell annotation tools for each dataset. The datasets are grouped by platform (CyTOF and FCM), indicated by the grey bars on top of the heatmap. The rows represent the tools, ordered by their overall mean performance, and the columns represent the datasets. Dataset labels include the dataset abbreviation (Table 1), tissue type, number of markers used (M), and number of annotated populations (P). Boxplots on the right side of the heatmap summarize the performance distribution of each tool across all datasets and cross-validation folds. Boxplots underneath the heatmap summarize the performance of the tools per dataset, illustrating dataset complexity (the Random baseline is excluded from the bottom panel). In all boxplots, the center line represents the median, box limits indicate the upper and lower quartiles, and whiskers extend to 1.5× the interquartile range; points beyond the whiskers represent outliers. Results denoted with an asterisk (*) indicate that the respective tool was trained using only 10% of the dataset.”

#### Overall Performance

The tools demonstrated varying performance across the eight datasets. CyAnno had the highest F1-scores on most datasets (0.76 to 0.98). DGCyTOF consistently achieved second highest F1-scores overall (0.75–0.98). CyTOF Linear Classifier and CyGATE achieved comparable performance on most datasets (0.72-0.94) and (0.71-0.97) respectively, with CyTOF Linear Classifier achieving the highest F1 score on HumanBoneMarrow_flow across all the tools. GateMeClass was only evaluated on a subset of datasets due to computational constraints, achieving F1 scores ranging from 0.49 to 0.81 for the E run and 0.32 to 0.82 for the V run. kNN performed comparably to CyTOF Linear Classifier on the CyTOF datasets and HumanBoneMarrow_flow, while only performing slightly poorer on the remaining FCM datasets. The random classifier achieved F1 scores below 0.16 across all datasets.

#### Dataset-Specific Performance

Performance varied both across and within the two platforms. Among the CyTOF datasets, the ChikVirusPBMC_Cyt dataset yielded the lowest F1 scores across nearly all tools. The bone marrow and stimulated blood datasets (MouseBoneMarrow_Cyt, StimBlood_Cyt, and HumanBoneMarrow_Cyt) yielded the highest F1 scores, while the CyTOF PBMC datasets (ChikVirusPBMC_Cyt and PBMC_Cyt) performed the worst. Among the FCM datasets, CyAnno and DGCyTOF achieved notably higher F1 scores on the PBMC_flow and covidPBMC_flow datasets compared to all other tools.

#### Performance Consistency

The distributions on the right-hand boxplots of Figure 2 show that CyAnno and DGCyTOF achieved the highest median F1 scores across all datasets. While their interquartile ranges were relatively tight, the notably lower performance on the ChikVirusPBMC_Cyt dataset increased their spread. CyTOF Linear Classifier also showed a narrow interquartile range, though at a lower overall performance level. CyGATE and KNN showed comparable median performance to CyTOF Linear Classifier but with wider spread across datasets. GateMeClass-E and GateMeClass-V showed the lowest median performance and the widest interquartile ranges, with GateMeClass-V in particular showing a long lower whisker. As expected, the random classifier served as a lower bound across all datasets.

#### Performance across cell populations

The distribution of F1 scores for the most and least prevalent cell populations across a selection of datasets is shown in Figure 3, while Supplementary Figure 1 provides the complete results across all datasets and cell populations.

**Fig. 3.**
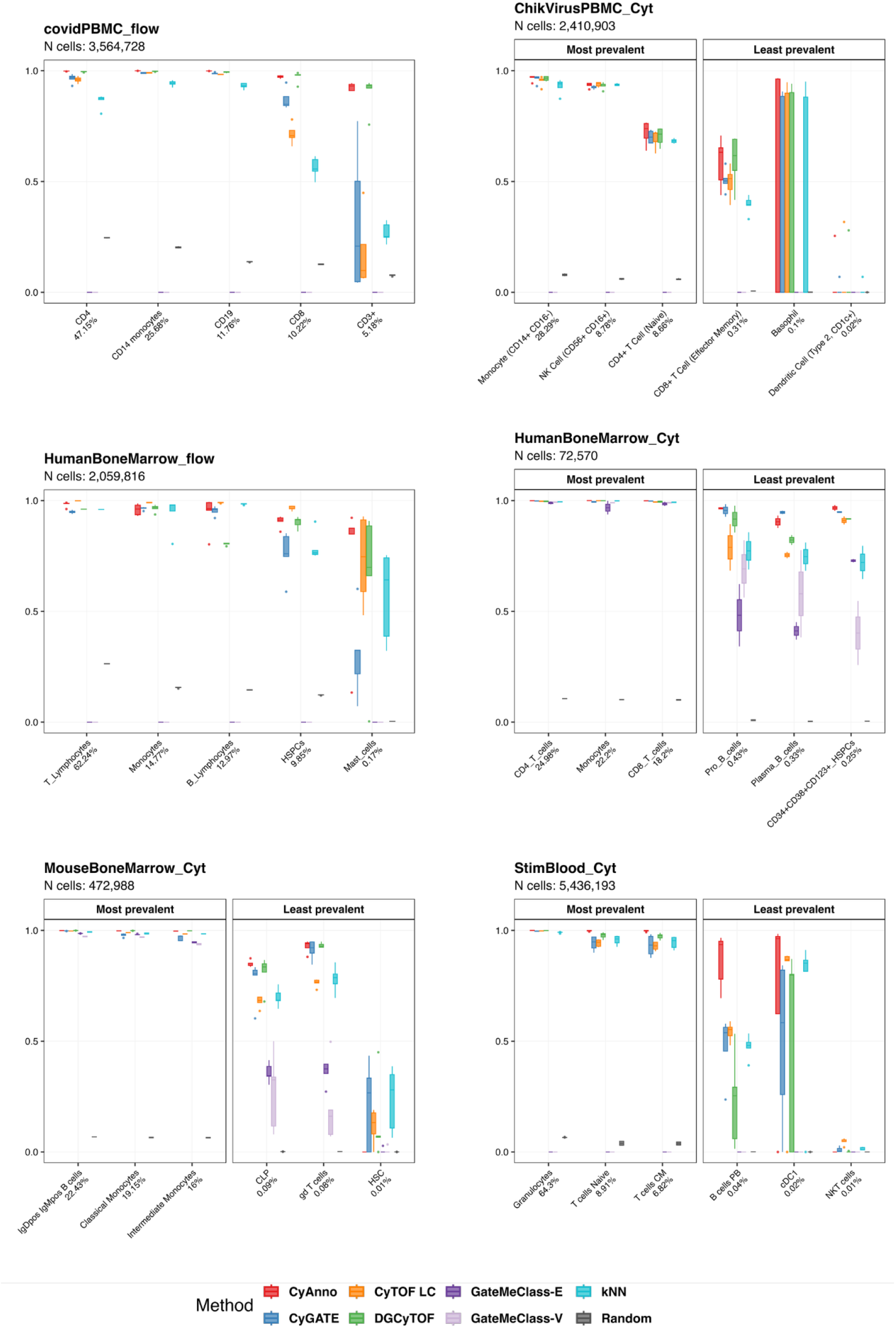
Performance of automated cell annotation tools on most and least prevalent cell populations across benchmark datasets. Boxplots illustrate the distribution of F1 scores for the top three most and least prevalent populations within six of the datasets. Populations are ordered by decreasing prevalent (left to right within each facet), with the specific cell type name and its percentage of the total population indicated on the x-axis. The total number of cells (N cells) for each dataset is stated in the subtitle. Boxplots represent the distribution of scores across cross-validation folds; the center line indicates the median, box limits represent the upper and lower quartiles, and whiskers extend to 1.5× the interquartile range. The colors correspond to the different annotation tools: CyAnno (red), CyGATE (blue), DGCytof (green), GateMeClass (purple), CyTOF Linear Classifier (orange), kNN (cyan), and Random (grey).

CyAnno showed high and consistent performance over cross-validation folds on both more and less prevalent cell types across all datasets, with the exception of the ChikVirusPBMC_Cyt dataset, where all tools showed reduced performance for less prevalent cell types. DGCyTOF performed well on more prevalent populations and maintained moderate performance on less prevalent ones across most datasets. CyGATE performed comparably to CyAnno and DGCyTOF on the more prevalent cell types in the covidPBMC_flow, HumanBoneMarrow_Cyt and MouseBoneMarrow_Cyt datasets, but was among the worst-performing tools on the StimBlood_Cyt and PBMC_Cyt datasets and on less prevalent cell types in the HumanBoneMarrow_flow dataset.

CyTOF Linear Classifier performed well on the HumanBoneMarrow_flow dataset and on the most prevalent cell types in the covidPBMC_flow, HumanBoneMarrow_Cyt and PBMC_Cyt datasets. Both CyGATE and CyTOF Linear Classifier showed high variance across cross-validation folds on some of the less prevalent cell types across datasets, indicating sensitivity to training data composition and poor generalization to unseen samples. GateMeClass achieved reasonable F1 scores on the most prevalent cell types in the HumanBoneMarrow_Cyt and MouseBoneMarrow_Cyt datasets, but performed considerably worse on less prevalent populations, with high variance across cross-validation folds further suggesting poor generalizability. GateMeClass performed poorly on the PBMC_flow dataset (Supplementary Figure 1), with some cross-validation runs performing worse than the random baseline. kNN generally performed well on the most prevalent cell types across dataset but was among the worst-performing tools on the less prevalent cell types.

#### Performance across dataset characteristics

To investigate the drivers of performance across datasets, we examined the relationship between mean F1 score and two dataset characteristics: number of markers and population (Supplementary Figure 2a,c) and mean cells per sample (Supplementary Figure 3a) . The performance fluctuated across both characteristics with no particular trend, suggesting that dataset size and dimensionality does not affect annotation accuracy.

Runtime was similarly unaffected by the both characteristics (Supplementary Figure 3b and Supplementary Figure 2b,d) with all tools showing broadly comparable runtime across datasets.

The effect of training size on both runtime and performance is shown in Supplementary Figure 3c-d. Most tools showed relatively stable runtimes across training sizes. CyAnno, DGCyTOF and kNN showed an uneven pattern, indicating the training size is not directly determining runtime. In terms of performance, both CyAnno and DGCyTOF showed notable F1 score improvement between 1,000 and 5,000 training cells, after which performance largely plateaued. CyAnno achieved the highest F1 scores across all tools. The remaining tools showed relatively flat performance trajectories with CyTOF Linear Classifier and kNN showing modest improvements with more training data but remained at moderate to low F1 scores, while CyGATE and GateMeClass showed inconsistent patterns across training sizes.

## Discussion

The presented benchmark evaluated the five described tools under a sample-centric cross-validation scenario intended to approximate a common clinical study scenario in which a single reference sample is used to annotate remaining samples. Across datasets, supervised approaches outperformed the random-label baseline, supporting that the standardized OmniBenchmark pipeline captures learnable structure aligned with the input gating labels. Within the benchmarked tools, CyAnno showed the most consistently high performance across datasets, with DGCyTOF typically close behind. CyTOF Linear Classifier and CyGATE displayed more context-sensitive behavior, and GateMeClass represents a constrained case because it was computationally intensive and could only be evaluated on a subset of datasets and under reduced training size, limiting direct comparability to methods trained on the full reference. More broadly, the pattern suggests that panel dimensionality and population separability interact with model capacity. In simpler settings, linear models can perform well, whereas in more complex datasets with heterogeneous cohorts and rare populations, robustness to imbalance and ambiguous population boundaries becomes more important.

Tools differed not only in peak accuracy but also in robustness to factors that commonly vary across studies including cohort heterogeneity and ambiguity near population boundaries. In this regard, CyAnno’s consistent strong performance under limited-reference settings is notable and is plausibly aided by its training strategy (kernel density estimation to identify high-confidence “core” cells before classification), which may reduce sensitivity to noise and transitional states. Conversely, approaches that rely on more global or geometry-sensitive assumptions can be more affected when populations are less prevalent or ambiguous.

Performance varied substantially across datasets, and this variation was not well explained by coarse descriptors such as cells per sample or marker-to-population ratio. Instead, differences likely reflect disease-associated distribution shifts, population separability, and annotation conventions, including how ungated populations are handled. Therefore, the same model can perform differently in canonical reference-like settings versus perturbed cohorts even when having overlapping markers. Accordingly, strong performance on widely used reference-style datasets should not be taken as evidence of broad generalizability. This implies that tool selection should ideally be validated on a representative subset of the target study rather than inferred from this benchmark.

Less prevalent populations represent a major practical bottleneck for automated annotation. In our benchmark, performance on low-prevalence classes (<1%) was less stable, consistent with sensitivity to limited examples and to which sample served as the reference. Methods also diverged most clearly in this area, with CyAnno generally retaining higher performance for less prevalent populations than several other approaches. These findings motivate two practical recommendations: (i) report per-population metrics alongside aggregate scores, and (ii) use targeted strategies to mitigate class imbalance (e.g., careful reference selection, or explicit unknown handling) when low-prevalence populations are biologically important.

Downsampling analyses further underscore the relevance of reference availability in real-world use. Performance improvements were concentrated at smaller reference sizes and then tended to plateau, suggesting no benefit from additional labeled cells beyond a modest reference set. At the same time, the relative stability of runtime with respect to training size suggests that collecting additional labeled reference data when feasible may be a more effective lever for improving performance than attempting to optimize computational throughput, particularly in clinically oriented settings where accuracy and robustness are paramount.

Several limitations should be acknowledged. First, the reliance on manual annotation to define ground truth inherently introduces operator dependence. Because manual gating can vary substantially between operators and centers, our evaluation effectively measures a model’s ability to replicate a specific manual annotation rather than an absolute biological truth. Consequently, models that correctly identify novel or overlooked signals (such as biologically relevant subsets hidden within “ungated” events) may be inadvertently penalized. Furthermore, additional work is needed to include multi-tube FCM datasets to capture performance across panels.

Second, methodological constraints shaped the scope of our analysis. To ensure objectivity and reproducibility, tools were strictly required to be completely autonomous (capable of training directly from labeled data without additional user decisions) and technically compatible with the Omnibenchmark framework. Consequently, several published annotation tools were excluded. DeepCyTOF [30], Cycadas [31], and CytoPheno [32] were omitted due to architectural incompatibilities with our automated pipeline or because of the use of outdated packages. Tools such as ACDC [33], and SCINA [34], were omitted based on the autonomy criteria as they require manual thresholds, intermediate clustering steps, or curated signatures. Attempting to develop automated wrappers to bypass these interactive or semi-supervised steps would constitute tool modification, fundamentally compromising the fairness and reproducibility of the benchmark. These mentioned tools could be further developed to be compatible with the omnibenchmark framework.

Finally, most tools were evaluated using default or minimally adjusted parameters rather than extensive dataset-specific hyperparameter optimization. This emphasizes out-of-the-box utility but may underestimate best-case performance for methods that benefit from careful tuning. Future work should expand benchmark coverage to parameter tuning, additional tissues and perturbations, evaluate alternative reference regimes (e.g., multi-donor references), and more systematically assess how preprocessing choices, data transformation, ungated-cell handling, and tuning strategies affect both overall performance and rare-population sensitivity.

From a practical standpoint, tool selection should be treated as a deployment decision. Users should validate candidate methods on a small representative subset of their own study using the intended preprocessing and labeling scheme, and inspect per-population performance rather than relying on a single aggregate score. When rare populations are of interest, reference selection and explicit handling of ambiguous or ungated events become particularly important, since these choices can dominate performance in low-prevalence settings.

## Conclusion

In this study, we present a comprehensive, open-source benchmark for automated cytometry annotation tools under a standardised, sample-centric cross-validation framework that reflects how these methods are typically deployed in practice. We evaluated robustness under constraints that frequently determine success in real projects, including rare-population detection, limited training data, and computational scalability.

By testing across a variety of datasets, we get a more general picture of performance of tools while the foundation for tool selection for individual use cases.

The reproducible and extensible benchmark enforces consistent data handling and evaluation across tools, providing a consistent method for evaluation. Importantly, this benchmark is designed as a living resource within the Omnibenchmark framework.

By providing a structured, transparent, and reproducible platform for researchers and developers, new tools can be rigorously performance tested to iteratively improve annotation algorithms in response to emerging biological and computational challenges. We invite the community to contribute datasets, tools, and refinements, ensuring that this benchmark remains a dynamic foundation for advancing automated cytometry analysis.

