## Supplementary Figures for "A Reproducible and Extensible Benchmark of Supervised Cell Type Annotation Tools for Cytometry Data"

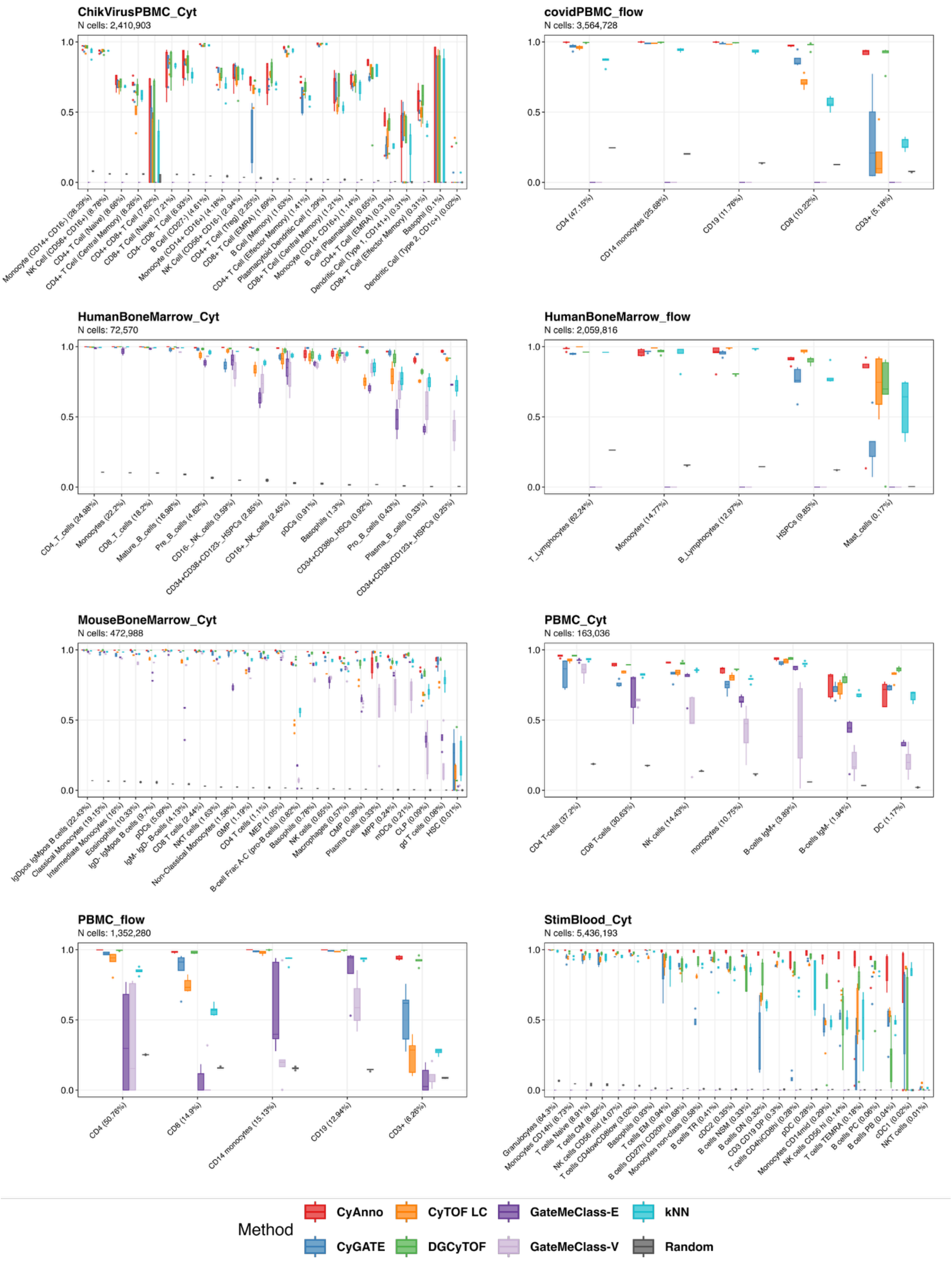


**Supp. Fig. 1 Performance of automated cell annotation tools across cell populations and benchmark datasets.** Boxplots illustrate the distribution of F1 scores for all the evaluated datasets. Populations are ordered by decreasing prevalence (left to right within each facet), with the specific cell type name and its percentage of the total population indicated on the x-axis. Boxplots represent the distribution of scores across cross-validation folds; the center line indicates the median, box limits represent the upper and lower quartiles, and whiskers extend to 1.5× the interquartile range. The colors correspond to the different annotation tools: CyAnno (red), CyGATE (blue), DGCytof (green), GateMeClass (purple), CyTOF Linear Classifier (orange), kNN (cyan), and Random (grey).

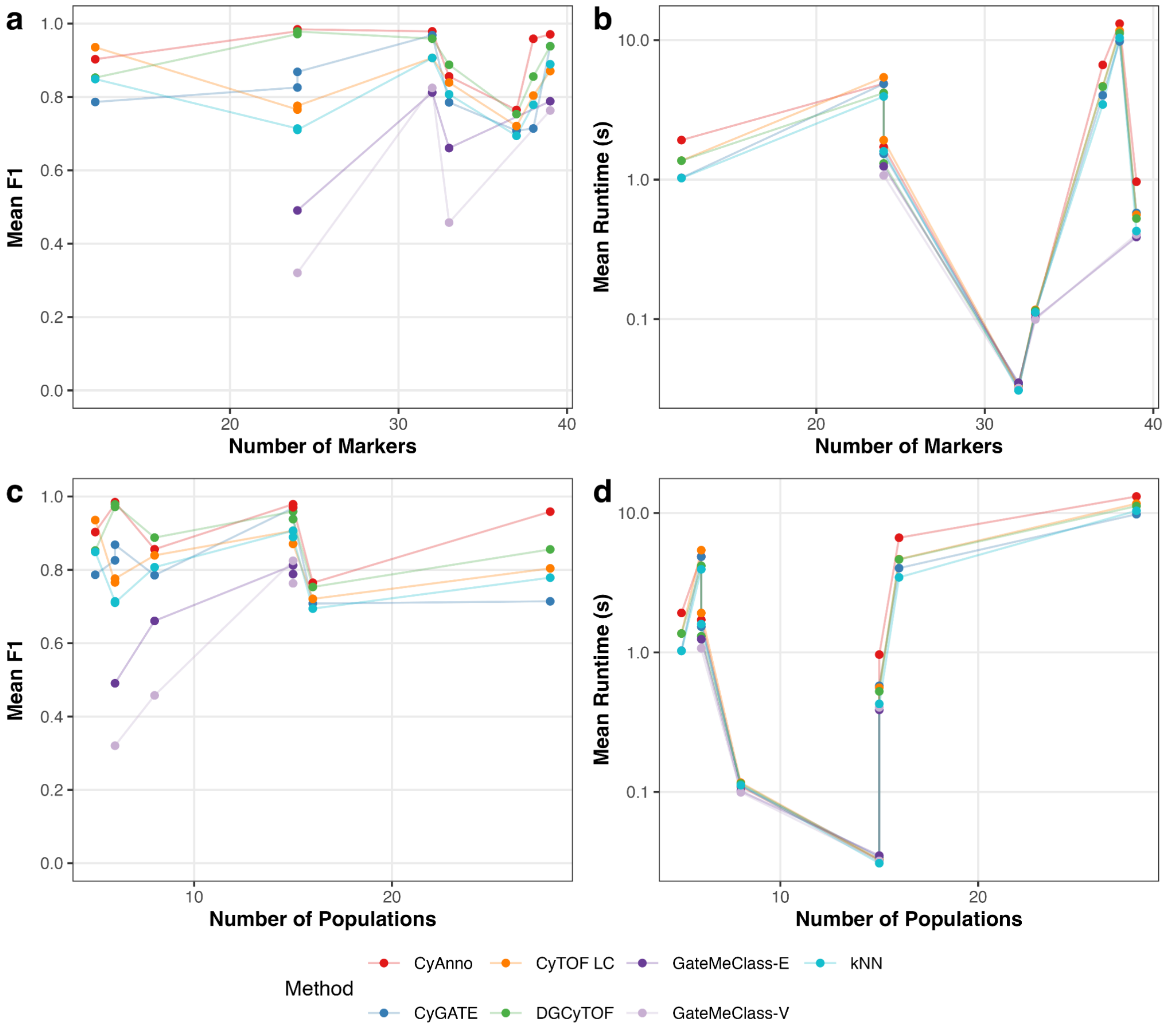


**Supp. Fig. 2 Impact of dataset dimensionality and population complexity on automated annotation tools.** Line charts illustrating how the number of markers and number of populations influence model performance and resource consumption. **a-b)** Relationship between the number of available markers and: **a)** classification performance (Mean F1-score) and **b)** mean runtime in seconds (log10 scale). **c-d)** Relationship between the total number of annotated cell populations and: **c)** classification performance (Mean F1-score) and **d)** mean runtime in seconds (log10 scale). Each point in panels a-d represents the mean performance or run time of a tool on a specific benchmark dataset.

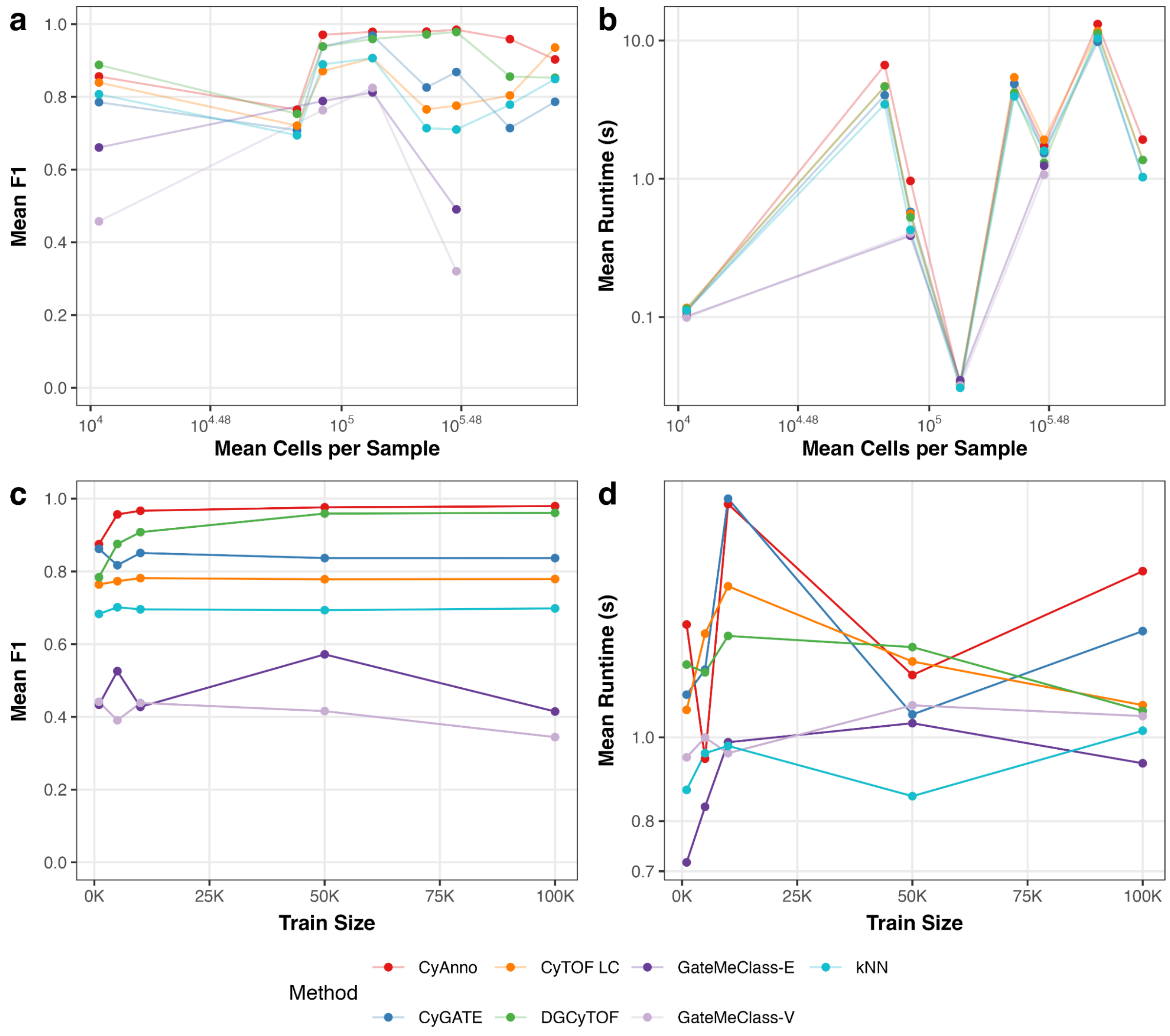


**Supp. Fig. 3 Impact of dataset size and training set size on performance and runtime.** Line charts illustrating how sample cell counts and training data volume impact model performance and computational efficiency. **a-b)** Relationship between dataset size (mean number of cells per sample, log10 scale) and: **a)** classification performance (Mean F1-score) and **b)** mean runtime in seconds (log10 scale). Each point in panels a-b represents the average performance of a tool on a specific benchmark dataset. **c-d)** Performance and scalability assessment using data subsampled to varying training sizes (ranging from 1k to 100k cells). **c)** Classification performance, showing the mean F1-score across increasing training volumes. **d)** Computational scalability, showing runtime (log10 scale) as a function of training set size.
